# The demographic costs of sexually antagonistic selection in partially selfing populations

**DOI:** 10.1101/2022.01.23.477381

**Authors:** Colin Olito, Charlotte de Vries

## Abstract

Classic population genetics theory has been fundamental to understanding the evolution of sex-differences and the maintenance of sexually antagonistic (SA) genetic variation, but these models have rarely considered the demographic consequences of intralocus sexual antagonism. In this paper we develop a stage-structured mendelian matrix model and jointly analyze the evolutionary and demographic consequences of SA selection in obligately outcrossing (i.e., dioecious/gonochorous) and partially selfing hermaphrodite populations. We focus on identifying parameter conditions under which SA polymorphism is maintained *and* the population growth rate remains positive. Additionally, we analyze the effects of inbreeding depression manifesting at different life-history stages and give an illustrative example of the potential for SA polymorphism in real populations using empirically estimated demographic rates for the hermaphroditic flowering plant *Mimulus guttatus*. Our results show that when population intrinsic growth rates approach one, extinction occurs across large swathes of parameter space favoring SA polymorphism or the fixation of male-beneficial alleles, and that inbreeding depression is a significant problem for maintaining SA polymorphism in partially selfing populations. Despite these demographic challenges, our example with *M. guttatus* appears to show that demographic rates observed in some real populations are capable of sustaining large regions of viable SA polymorphic space.

## Introduction

To persist in the long term, a population must be able to adapt to its environment. Yet, the process of adaptation can be impeded by a variety of genetic and environmental factors, such as deleterious mutations (Haldane, 1937), changing environmental conditions (Lande and Shannon, 1996; Maynard Smith, 1976; Orr and Unckless, 2008), maladaptive gene-flow (Bolnick and Nosil, 2007; Kirkpatrick and Barton, 1997), and genetic constraints (Connallon and Hall, 2018; Matthews et al., 2019). Genetic constraints on adaptation can arise as a consequence of conflicting selection and gene-flow between different classes of individuals, or more generally, when there are genetic trade-offs between fitness components (Charlesworth and Hughes, 2000; Connallon and Hall, 2018).

Fitness trade-offs that impose genetic constraints on adaptation have particularly interesting evolutionary consequences. On the one hand, they prevent individuals (or classes of individuals) from reaching their phenotypic optimum in one or more fitness components, which can increase a population’s overall extinction risk (Harts et al., 2014; Kokko and Brooks, 2003). On the other hand, they provide an effective mechanism for the maintenance of genetic variation, which can increase a population’s capacity for future adaptation (Charlesworth and Hughes, 2000; Connallon and Hall, 2018; Fisher, 1930; Matthews et al., 2019). Hence, for traits under conflicting selection, the nature and extent of genetic variation observed in natural populations should reflect a balance between the maintainance of genetic polymorphisms, and the population dynamical consequences of the resulting maladaptation.

Sexually antagonistic selection (abbreviated SA hereafter) arising from genetic trade-offs between male and female fitness, is a common feature of sexually reproducing populations, and is thought to contribute to the maintenance of genetic variation and the evolution of sexual dimorphism and sex-related traits (Bonduriansky and Chenoweth, 2009; Charlesworth, 1999; Lande, 1980; Olito and Connallon, 2019; Rice, 1992; Rice and Chippindale, 2001). When such genetic trade-offs occur, SA selection can arise when beneficial alleles for one sex are deleterious when expressed in the other (Connallon and Clark, 2012; Kidwell et al., 1977; Rice, 1992).

Alleles with opposing fitness effects through male and female sex functions can cause analogous genetic constraints on fitness in hermaphrodite populations, where both maternal and paternal reproductive success contribute jointly to each individuals’ overall fitness (Abbott, 2011; Jordan and Connallon, 2014; Lloyd and Webb, 1986; Webb and Lloyd, 1986). Self-fertilization in hermaphrodites is predicted to reduce the total parameter space where balanced SA polymorphisms can be maintained, while simultaneously creating a bias in selection through the female sex function (Glémin 2021; Jordan and Connallon 2014; but see Tazzyman and Abbott 2015). Yet, other factors, such as genetic linkage to other SA loci or a sex-determining region (Jordan and Charlesworth, 2012; Olito, 2017; Olito and Connallon, 2019; Otto et al., 2011), antagonism between viability and fecundity selection (Glémin, 2021), or spatial heterogeneity and complexity of the life-cycle can expand the parameter space for SA polymorphism (Connallon et al., 2019; Glémin, 2021; Olito et al., 2018). Overall, both theoretical predictions and current empirical data suggest that there is ample scope for SA trade-offs and the maintenance of SA polymorphisms in both dioecious and hermaphrodite populations (Abbott, 2011; Wang et al., 2020), although identifying specific SA loci from genome sequence data remains challenging (Ruzicka et al., 2020).

Most population genetic models, including models of the maintenance of SA polymorphisms, keep track of the relative fitnesses of genotypes in populations of constant (and often infinite) size, and therefore do not consider the population dynamical consequences of evolutionary change. In reality, these genotypic fitnesses emerge from myriad processes acting throughout the life-history of individuals (e.g., Johnston et al. 2013; Mérot et al. 2020). To take into account the consequences of population dynamics under SA selection, a model is required that links changes in genotype distributions to population dynamics. The potential demographic costs of sexual antagonism were pointed out by Kokko and Brooks (2003), but few papers since then have explicitly incorporated demography into models of sexual antagonism (with the notable exception of Harts et al. 2014). de Vries and Caswell (2019a) introduced a mendelian matrix model with intralocus sexual antagonism, and population dynamics but did not perform an analysis of the consequences of demographic viability for the scope of sexual antagonism to maintain polymorphism.

In this paper, we extend the model of (de Vries and Caswell, 2019a,b) to include both dioecious and hermaphroditic species in order to study the parameter conditions under which different outcomes of SA selection, including fixation of male- or female-beneficial alleles as well as balanced polymorphisms, are also demographically viable. A major strength of our approach is that the model can be parameterized using empirically estimated demographic rates, enabling us to make predictions about the scope for demographically viable SA polymorphisms that are grounded in the biology of real populations. We demonstrate this with a case study of *Mimulus guttatus* (now *Erythranthe guttata*)

### The Model

Here, we briefly describe a matrix model incorporating multiple life-cycle stages and a single diallelic locus under SA selection on male and female fertility for populations of partially-selfing simultaneous hermaphrodites. The derivation and some of the key results follow closely those presented in de Vries and Caswell (2019a) and de Vries and Caswell (2019b). In fact, the model reduces to the stage-structured model of de Vries and Caswell (2019a) under obligate outcrossing. A full derivation of the model and analyses is presented in the Online Supplementary Material, and all computer code necessary to reproduce the results are available at https://anonymous.4open.science/r/SA-Hermaphrodites-wDemography-BF66.

Simultaneous hermaphrodites can transmit genes to the next generation via both sperm/pollen and eggs/ovules, and have the potential to reproduce by a combination of self- and outcross-fertilizations. Maternal outcrossing involves receiving male gametes from another individual in the population, while paternal outcrossing involves exporting male gametes to another individual and fertilizing their ovules. Self-fertilization is achieved when an individual’s male gametes fertilize their own ovules. To distinguish between parameters relating to male and female function, we denote matrices or vectors relating to the male sex function with a prime.

Individuals in the model are jointly classified by life-cycle stage (1, …, *w*), genotype (1, …, *g*), and whether they were produced by self- or outcross fertilization (denoted by *S* and *X* superscripts, respectively; see Table 1 for a full description of terms included in the model). Whether individuals were produced through selfing or outcrossing is included in the individual state description because individuals produced through selfing might experience reduced survival, growth, or maturation rates as a consequence of inbreeding depression. In reality, the severity of inbreeding depression will not only be a function of whether an individual is produced by selfing but also of how many consecutive generations of inbreeding have occurred in their lineage (e.g., Kelly 1999, 2007). However, tracking lineage-specific inbreeding histories is beyond the scope of this paper, and we therefore make the common assumption that the severity of inbreeding depression is the same for all individuals (e.g., Charlesworth and Charlesworth 1987, 2010; Charlesworth and Willis 2009; Jordan and Connallon 2014).

**Table 1:**
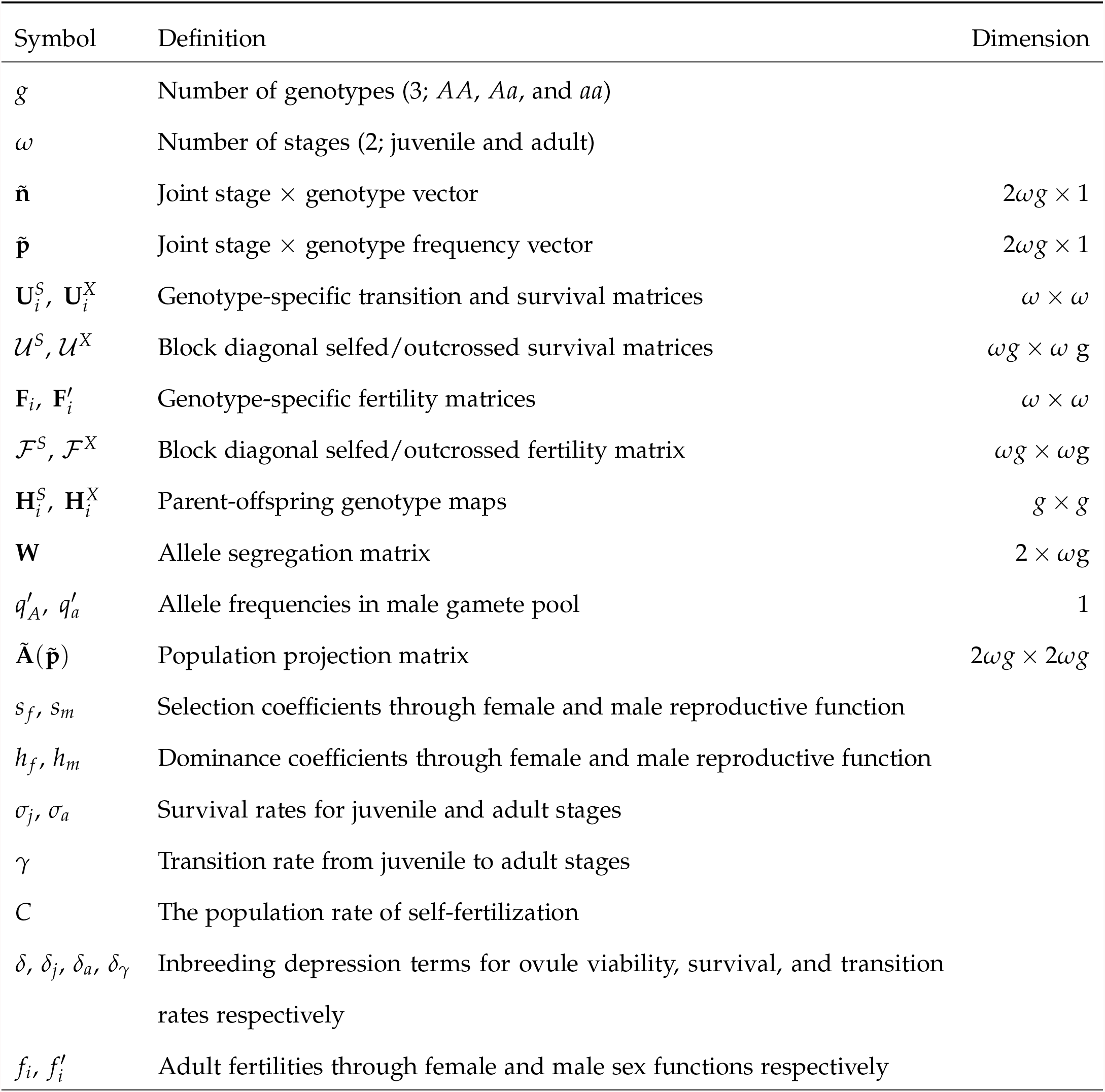
Definition of terms.

| Symbol | Definition | Dimension |
| --- | --- | --- |
| $g$ | Number of genotypes (3; $AA$, $Aa$, and $aa$) | |
| $\omega$ | Number of stages (2; juvenile and adult) | |
| $\tilde{\mathbf{n}}$ | Joint stage × genotype vector | $2\omega g \times 1$ |
| $\tilde{\mathbf{p}}$ | Joint stage × genotype frequency vector | $2\omega g \times 1$ |
| $\mathbf{U}_i^S, \mathbf{U}_i^X$ | Genotype-specific transition and survival matrices | $\omega \times \omega$ |
| $\mathcal{U}^S, \mathcal{U}^X$ | Block diagonal selfed/outcrossed survival matrices | $\omega g \times \omega g$ |
| $\mathbf{F}_i, \mathbf{F}'_i$ | Genotype-specific fertility matrices | $\omega \times \omega$ |
| $\mathcal{F}^S, \mathcal{F}^X$ | Block diagonal selfed/outcrossed fertility matrix | $\omega g \times \omega g$ |
| $\mathbf{H}_i^S, \mathbf{H}_i^X$ | Parent-offspring genotype maps | $g \times g$ |
| $\mathbf{W}$ | Allele segregation matrix | $2 \times \omega g$ |
| $q'_A, q'_a$ | Allele frequencies in male gamete pool | 1 |
| $\tilde{\mathbf{A}}(\tilde{\mathbf{p}})$ | Population projection matrix | $2\omega g \times 2\omega g$ |
| $s_f, s_m$ | Selection coefficients through female and male reproductive function | |
| $h_f, h_m$ | Dominance coefficients through female and male reproductive function | |
| $\sigma_j, \sigma_a$ | Survival rates for juvenile and adult stages | |
| $\gamma$ | Transition rate from juvenile to adult stages | |
| $C$ | The population rate of self-fertilization | |
| $\delta, \delta_j, \delta_a, \delta_\gamma$ | Inbreeding depression terms for ovule viability, survival, and transition rates respectively | |
| $f_i, f'_i$ | Adult fertilities through female and male sex functions respectively | |

The population state at time *t* is described by a population state vector, **ñ**(*t*), which is ordered by how individuals were produced (selfing vs. outcrossing), then by genotype, and finally by stage. For a single diallelic locus with alleles *A* and *a*, we have three genotypes (*AA, Aa, aa*; *g* = 3), giving the population state vector:

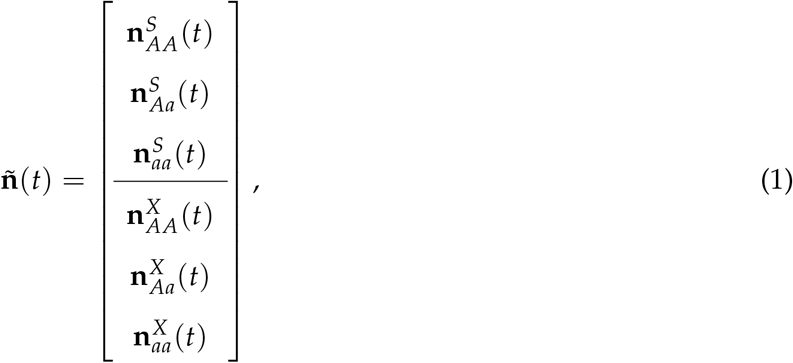

where 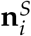 and 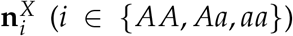 are the stage distribution vectors of individuals of genotype *i* produced by self-fertilization and outcrossing, respectively. The proportional population vector is given by

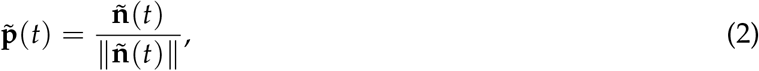

where ‖ ·‖ is the one-norm. The population vector **ñ**(*t*) is projected forward from time *t* to *t* + 1 by the projection matrix **Ã**(**ñ**) such that

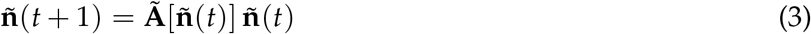

The population projection matrix **Ã** is constructed from four sets of matrices representing the demographic and genetic processes: The matrices 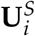 and 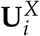 contain transition and survival probabilities for each genotype, produced by selfing and outcrossing respectively. The matrices **F**_*i*_ and 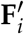 contain the genotype × stage specific contribution of genotype *i* to the female and male gamete pools, respectively, and therefore to zygotes in the next generation. We assume that whether individuals were produced through selfing or outcrossing does not affect their fecundity or mating success, that is, we assume 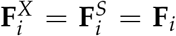 and 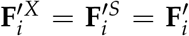. Deviations from this assumption are straightforward to incorporate but beyond the scope of this paper.

For the purpose of this article, we assume that mating is random with respect to stage and hence that the parent-offspring map is the same for all stages (i.e., 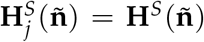, and 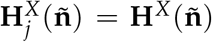; where *j* ∈ {1, *w*}). The matrices **H**^*S*^(**ñ**) and **H**^*X*^(**ñ**) contain the population genetic processes and are presented in the next section.

#### Mating and offspring production under partial selfing

Outcrossing in our model proceeds similarly to a two-sex model of reproduction (de Vries and Caswell, 2019b). That is, each individual’s genotype determines both the number of ovules produced and the paternal mating success, broadly defined. For hermaphroditic flowering plants, for example, paternal mating success could reflect pollen production, export efficiency, pollen-tube germination and growth rates, among other things (Harder et al., 2016; Lloyd and Webb, 1986; Wang et al., 2020). Note, however, that by modeling paternal relative mating success rather than pollen/sperm production, we implicitly assume female demographic dominance (i.e., production/transport of male gametes does not limit ovule fertilization and hence population growth), a point we return to in the Discussion.

Because we assume that inbreeding depression only affects offspring viability, the allele frequency in the male mating population is obtained by simply summing the vectors of individuals produced through selfing and outcrossing:

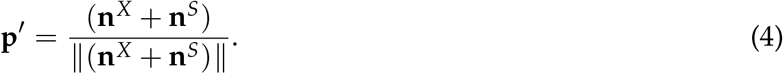

Depending on their stage and genotype at the SA locus, individuals contribute male gametes to the overall population gamete pool according to the following equation

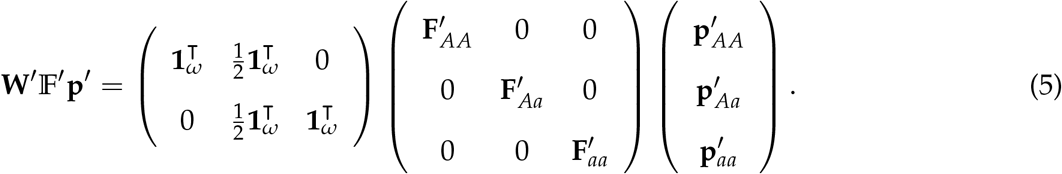

The matrix 𝔽′ is composed of genotype-specific fertility matrices, and operates on the the vector of genotype frequencies to give their relative contribution of each genotype to the gamete pool. The matrix **W**′ converts these contributions to allele numbers. Normalizing the resulting vector gives the allele frequencies in the male gamete pool:

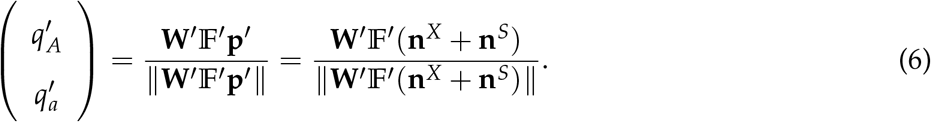

The key difference between selfing and outcrossing reproduction can be seen in the parent-offspring maps, which reflect the joint effects of meiosis and mating on the distribution of offspring genotypes.

The parent-offspring map for outcrossing is a function of the allele frequencies in the male gamete pool, and is given by

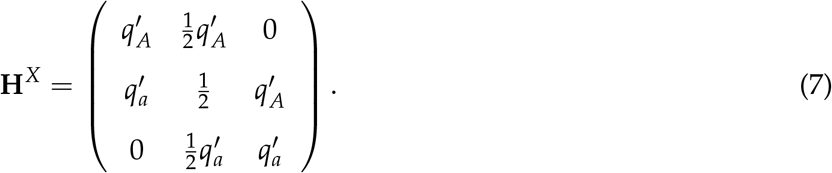

From left to right, the columns of matrix **H**^*X*^ give the genotype distribution of outcrossed offspring produced by a maternal parent of each genotype (*AA, Aa*, and *aa* respectively).

The parent-offspring map for reproduction via self-fertilization differs from the outcrossing parent-offspring map. As in previous models of partial selfing, we assume that individuals produce enough male gametes to easily self-fertilize all of their ovules, and that self-fertilization involves little or no selection via the male sex function from external factors (e.g., pollinator visitation) relative to outcrossing (Charlesworth and Charlesworth 1978; Jordan and Connallon 2014; Olito 2017, but see Tazzyman and Abbott 2015). Under these assumptions, the genotype distributions of selfed offspring are determined entirely by the parental genotype and the probabilities of segregation and fertilization during and after meiosis:

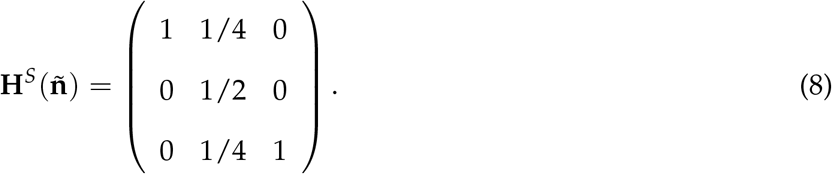

The columns of **H**^*S*^(**ñ**) again give the selfed offspring genotype distributions for parental genotypes of *AA, Aa*, and *aa*, respectively. As described in the next section, these parent-offspring maps influence the projection matrix by altering the fertilitiy matrices for individuals produced by selfing and outcrossing (see Eq(12) and Eq(13) below).

Following previous theory for partially selfing populations (e.g., Charlesworth and Charlesworth, 2010; Glémin, 2021; Jordan and Connallon, 2014), we use a ‘fixed prior selfing’ model where all individuals are assumed to self-fertilize at a constant rate, *C*, prior to receiving outcross pollen, for reasons of analytic tractability. In the Online Supplement, we present a more general model of genotype-specific self-fertilization rates (after Jordan and Connallon 2014).

#### Population projection

Using the component matrices described above (the survival matrices, 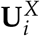,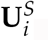, the fertility and mating success matrices, 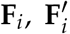, and the parent-offspring matrices **H**^*S*^(**ñ**) and **H**^*X*^), we construct the population projection matrix **Ã**[**ñ**] using the vec-permutation approach of Caswell et al. (2018), see Online Supplementary Material for the step by step construction of the model. The matrix that projects the eco-evolutionary dynamics is:

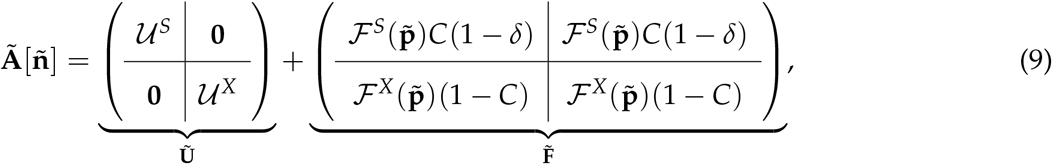

where *C* denotes the proportion of each individual’s ovules that are self-fertilized (the remaining 1 −*C* are outcrossed), and *δ* represents the proportion of self-fertilized zygotes that fail to develop due to inbreeding depression during early development (Charlesworth and Charlesworth, 1987).

The blocks of the component matrices in Eq(9) correspond to production of offspring by self-fertilization and outcrossing (ℱ ^*S*^ and ℱ ^*X*^ in 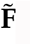), and survival of extant individuals produced by selfing or outcrossing (𝒰^*S*^ and 𝒰^*X*^ in **Ũ**). The survival matrices for individuals produced through selfing and outcrossing are,

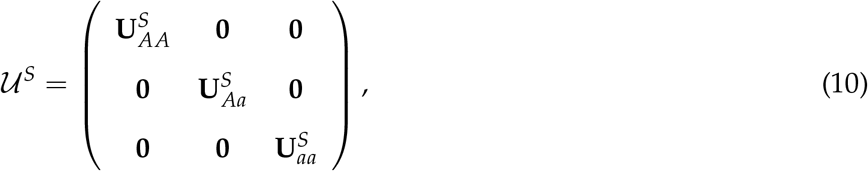

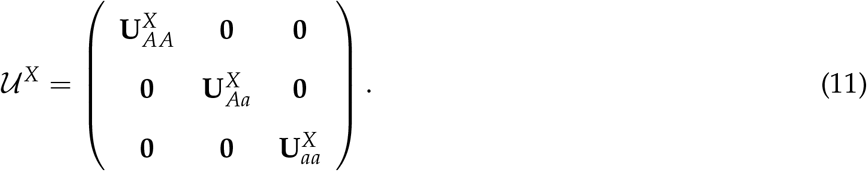

Since individuals cannot change their genotype once they are born, the survival matrices are block diagonal. Similarly, we construct fertility matrices for individuals produced through selfing and outcrossing,

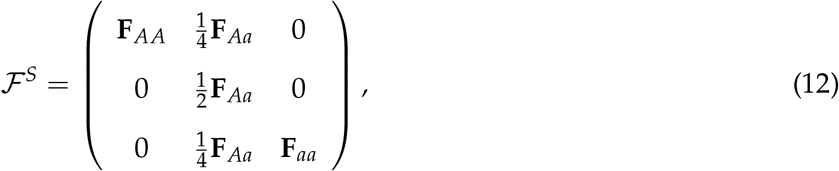

and

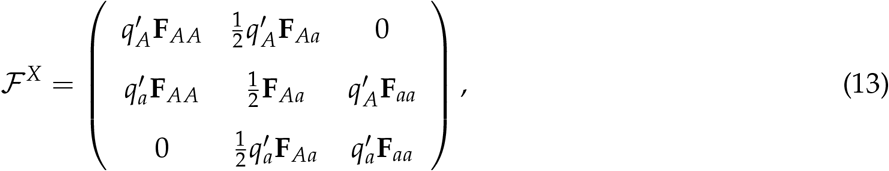

where 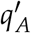 and 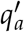 are given by Eq(6).

The blocks of 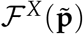 can be constructed and interpreted as follows: The first row block of the first column produces *AA* offspring by outcrossing from *AA* maternal parents. This happens when the *AA* maternal parent receives an *A* gamete from the male gamete pool, which happens with probability 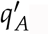. The other blocks can be interpreted similarly.

Combining all the component matrices yields the overall eco-evolutionary projection matrix shown in Appendix A.

#### Sexually antagonistic selection & inbreeding depression

We now construct and analyze a genotype × stage-classified model for a hypothetical species with intralocus sexual conflict via the two sex functions. For the sake of simplicity, we assume our hypothetical species has a life cycle with only two stages: juveniles and adults (i.e., *w* = 2), and that only adults are reproductively active. Suppose that there is a genetic trade-off between the sex-functions at a single diallelic locus such that allele *A* is beneficial for female fertility but detrimental for male reproductive success (e.g., pollen production), and that allele *a* has the reverse effect. Following convention, we parameterize the fertility component of fitness for each genotype through each sex function, *w*_*i*_ and 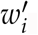, to be bounded by [0, 1], with dominance and selection coefficients *h*_*f*_, *s*_*f*_ and *h*_*m*_, *s*_*m*_ determining the decrease in fertility through each sex function relative to the most fit genotype (*AA* has highest female fertility, *aa* the highest male fertility; see Table 2).

**Table 2:** Relative fertilities for Sexually Antagonistic selection (*w*_*i*_)

|  | *Genotype* |  |  |
| --- | --- | --- | --- |
|  | *AA* | *Aa* | *aa* |
| Female function ($w_i$): | 1 | $1 - h_f s_f$ | $1 - s_f$ |
| Male function ($w'_i$): | $1 - s_m$ | $1 - h_m s_m$ | 1 |

The SA locus does not affect survival or transition rates. However, the survival matrices can be used to model the fitness effects of inbreeding depression at later stages of development by allowing different stage-specific survival and transition rates for individuals produced by self-fertilization vs. outcrossing. By contrast, the parameter *d* only affects inbreeding depression through viability of selfed ovules. With this in mind, we define survival matrices for individuals produced by selfing and outcrossing as follows:

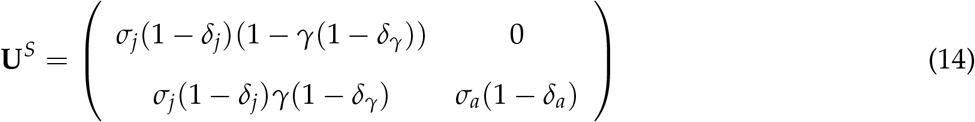

and

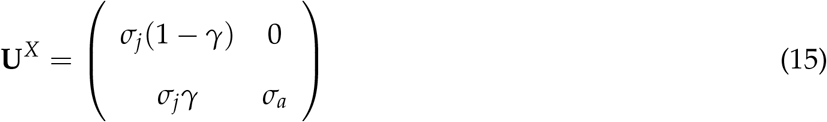

where *σ*_*j*_ and *σ*_*a*_ are the juvenile and adult stage survival rates, *γ* is the maturation rate from juvenile to adult stages, and the corresponding *δ*_*j*_, *δ*_*a*_, and *δ*_*γ*_ terms denote the proportional decreases in stage-specific survival and transition rates due to inbreeding depression (i.e., deleterious effects of inbreeding at later life-history stages; e.g., Harder and Routely 2006). For simplicity, we assume survival and transition rates are constant among genotypes.

Throughout our analyses, we distinguish between early- and late-acting inbreeding depression. We quantify early-acting inbreeding depression using *d*, and late-acting inbreeding depression using *δ*_*i*_ (where *i*∈ {*j, a*, γ})·*δ* denotes the fraction of self-fertilized ovules that do not develop into juveniles due to inbreeding depression. An important difference between early- and late-acting inbreeding depression in the model is that *δ*, affects the production of new individuals, whereas the *δ*_*i*_ affect the demographic rates of extant individuals, contained in **U**^*S*^ (see Eq. 14).

The fertility matrices through female and male function are

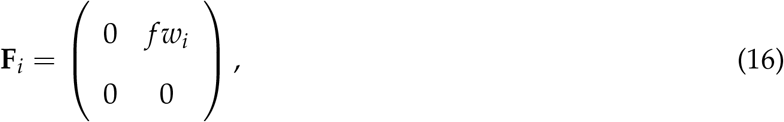

and

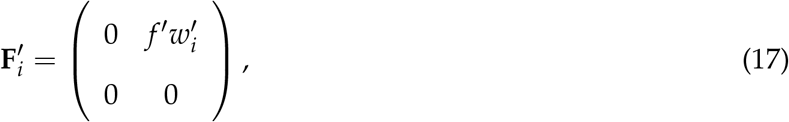

where *f* and *f* ′ represent adult fertilities, and *w*_*i*_ and 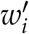 the genotypic relative scaling factors for female and male sex-functions (see Table 2).

Iterating the projection matrix, Eq(9) with the above demographic matrices, given an initial population state vector, allows numerical simulation of the eco-evolutionary dynamics for selection operating on any of the sex-function, or stage-specific demographic parameters. As we outline below, we use numerical techniques together with mathematical analyses to study the conditions for the maintenance of SA polymorphisms, and the demographic fate of the populations (i.e., positive growth, or extinction).

Unless stated otherwise, we use the following parameter values for the demographic rates in the model: *σ*_*j*_ = *σ*_*a*_ = 0.6 and *γ* = 0.05. These chosen values are similar to those used in de Vries and Caswell (2019b), facilitating comparison between models, and correspond to a life-history in which individuals spend multiple timesteps in the juvenile phase prior to maturing into reproductively active adults, but are otherwise arbitrary. Our parameters of interest include fertility, *f*, the inbreeding depression parameters, *δ, δ*_*j*_, *δ*_*a*_, *δ* _*γ*_, and the selection parameters *h* _*f*_, *s* _*f*_, *h*_*m*_, and *s*_*m*_, which are given different values for each analyses as described in the figure captions.

#### Analyses

Diverse eco-evolutionary outcomes are possible in the model, including fixation of either allele, balanced polymorphism, population growth or extinction, and even evolutionary rescue and suicide (e.g., see de Vries and Caswell 2019a,b). We focus on identifying parameter conditions where two criteria are satisfied: (*i*) SA polymorphism is maintained under balancing selection and (*ii*) the intrinsic population growth rate at equilibrium is positive; a situation that we refer to as a ‘demographically viable SA polymorphism’.

We identify conditions where SA polymorphism is ‘protected’ by evaluating the stability of populations initially fixed for either SA allele to invasion by the other (i.e., we assessed stability at the boundary equilibrium genotype frequencies of 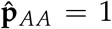 and 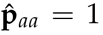; de Vries and Caswell 2019b; Levene 1953; Prout 1968). The formal conditions for a protected polymorphism are determined by linearizing the model in the vicinity of the boundary equilibria 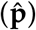, and evaluating the magnitude of the largest eigenvalue of the Jacobian matrix of the linearization. A full derivation of the Jacobian and details of the invasion analysis are provided in the Online Supplementary Material, and the relevant leading eigenvalues are presented in Appendix B.

We used numerical simulation to determine whether a protected SA polymorphism was also demographically viable. Specifically, for each boundary equilibrium we introduced the rare allele at low initial frequency, and iterated Eq(9) until the population had reached demographic and genotypic equilibrium. Our model implements density independent demographic rates, and so the population state vector will grow or shrink exponentially after converging to stable population structure and genotypic frequencies (see chapter 17 in Caswell 2001). The intrinsic population growth rate after convergence, *λ*, can be calculated as **ñ**(*t*)/**ñ**(*t* −1). We note, however, that if the ecological component of the model is non-linear, more exotic dynamics are possible (de Vries et al., 2020).

Because single-locus selection coefficients are generally weak (e.g., Eyre-Walker and Keightly 2007) and strongly skewed, we limit our analyses to coefficients within 0 < *s*_*f*_, *s*_*m*_ ≤ 0.15, unless stated otherwise. We present scenarios of equal dominance in the main text (i.e., *h*_*f*_ = *h*_*m*_ = *h*). Specifically, we examine scenarios of (*i*) additive SA fitness effects (*h* = 1/2), which are commonly observed for small-effect alleles or quantitative traits (Agrawal and Whitlock, 2011); and (*ii*) dominance reversals, where the deleterious fitness effect of each SA allele is partially recessive in each sex (*h* = 1/4), which are predicted under fitness landscape models provided the population is not too far from the phenotypic optimum (Connallon and Clark, 2014; Manna et al., 2011). We briefly explore the consequences of sex-specific dominance in the Online Supplementary Material.

For analyses of the demographic effects of inbreeding depression, we make two additional simplifying assumptions: First, to keep our analyses tractable we explore the effects of individual inbreeding depression terms in isolation. That is, we assume that only one of the *δ* and *δ*_*i*_ terms (where *i* ∈ {*j, a, γ*,}) can be non-zero at a time. Second, we assume that if inbreeding depression is caused primarily by recessive deleterious mutations (as suggested by empirical data), it should covary negatively with the population selfing rate due to purging, provided the population selfing rate has been relatively constant in recent evolutionary time (Charlesworth and Willis 2009; though we note that other processes could give rise to this pattern, e.g., Charlesworth and Willis 2009; Crnokrak and Barrett 2002; Hedrick and Garcia-Dorado 2016). Following Olito and Connallon (2019), we incorporate such negative covariance by constraining the inbreeding depression terms in the model (*δ* and *δ*_*i*_, where *i* ∈ {*j, a, γ*}) to follow a simple declining function of the selfing rate: *δ* = *δ*\*(1 −*b*(1 −*L*)), where *δ*\* is the hypothetical severity of inbreeding depression for a completely outcrossing population, *b* is a shape parameter determining how far *δ* will decline under complete selfing (when *C* = 1), and *L* describes the expected deleterious mutation load as a function of the selfing rate *C*. The function *L* includes an additional shape parameter, *a*, which determines the curvature of the overall function for *d* (see Appendix E in Olito and Connallon 2019 for additional details). We set *δ*\* = 0.8, *b* = 0.5, and *a* = 0.2 for all analyses, values chosen to be consistent with empirical estimates of inbreeding depression (e.g., fig. 2 in Husband and Schemske 1996).

## Results

### Fixation, Polymorphism, and Extinction

We begin with an illustration of demographically viable polymorphic parameter space in the absence of inbreeding depression in Figure 1 (i.e., *δ* = *δ*_*i*_ = 0). The red regions in Figure 1 indicate areas where the population goes extinct (population growth rate is less than one) for different values of the fertility parameter. Lower female fertility values correspond to a larger demographically inviable area. The funnel between the thick black lines indicates the area where the sexually antagostic alleles coexist. Above the funnel, the female-benefit and male-detriment allele goes to fixation; below the funnel, the male-benefit and female-detriment allele fixates. Much of the region where the male-benefit and female-detriment allele fixates is demographically inviable due to the demographic consequences of reduced female fitness in this region. The SA polymorphisms that remain viable at lower fertility values correspond to regions where the female-deleterious allele is predicted to segregate at low frequencies, or in regions with weak selection through both sex functions, an asymmetry that reflects the assumption of female demographic dominance.

**Figure 1:**
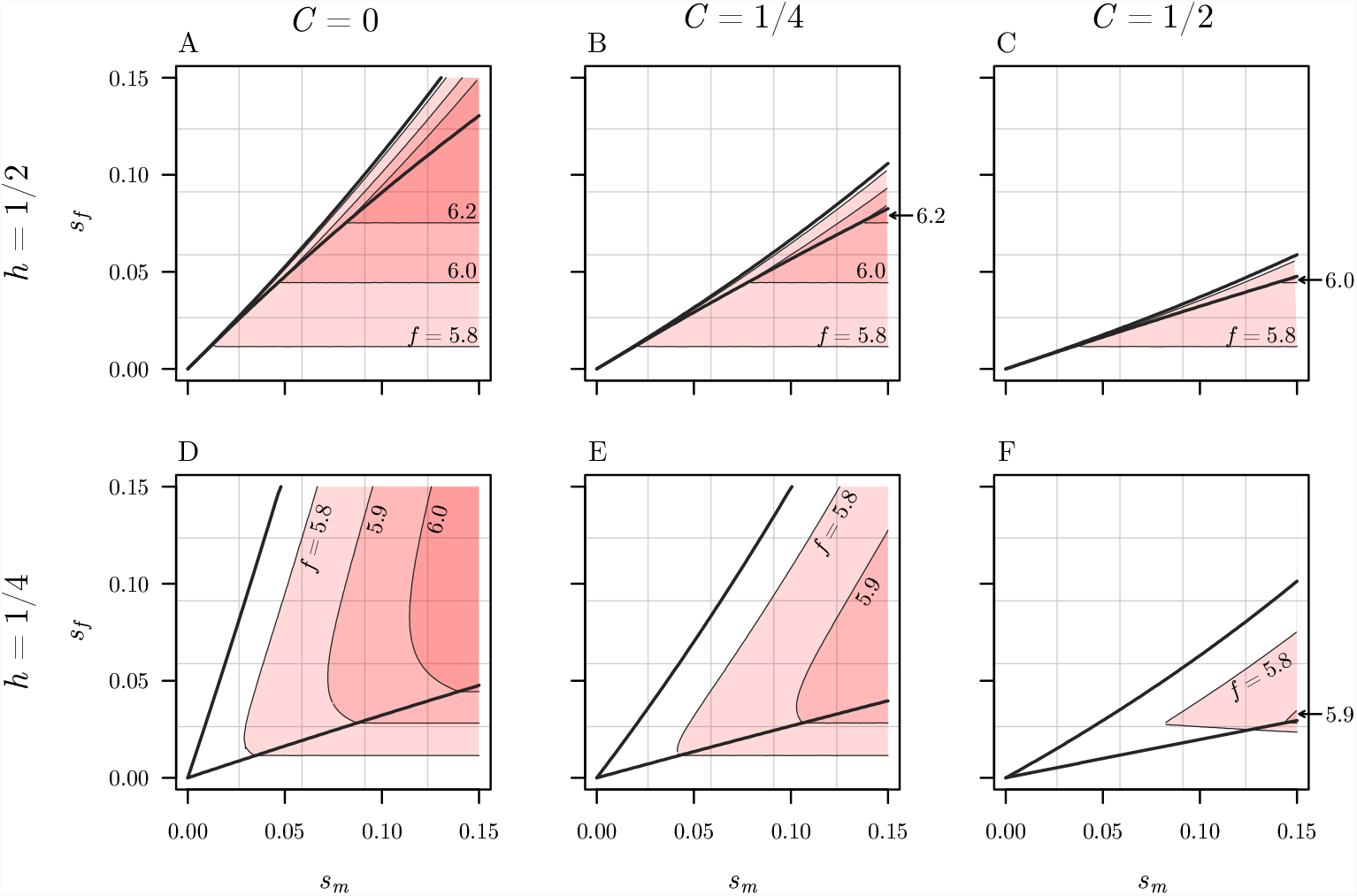
Illustration of parameter space for SA polymorphism and extinction thresholds predicted by the model. Balanced SA polymorphisms can be maintained in the funnel-shaped region between the invasion conditions for each SA allele (dark solid lines). However, for some parameter conditions, populations will ultimately go extinct (red shaded regions) due to reduced female fitness resulting from the male-beneficial/female-deleterious allele that is either segregating as a balanced polymorphism (inside the funnel), or becomes fixed (area below the funnel). “Demographically viable polymorphic parameter space” corresponds to the area inside the funnel that is also to the left of the extinction threshold for a given fertility value. Results are shown for three different population selfing rates (*C* = {0, 1/4, 1/2}), and two dominance scenarios (additivity, where *h* = 1/2, and dominance reversal, where *h* = 1/4); extinction thresholds are illustrated for three different values of female fecundity (*f* values annotated on each panel).

**Figure 2:**
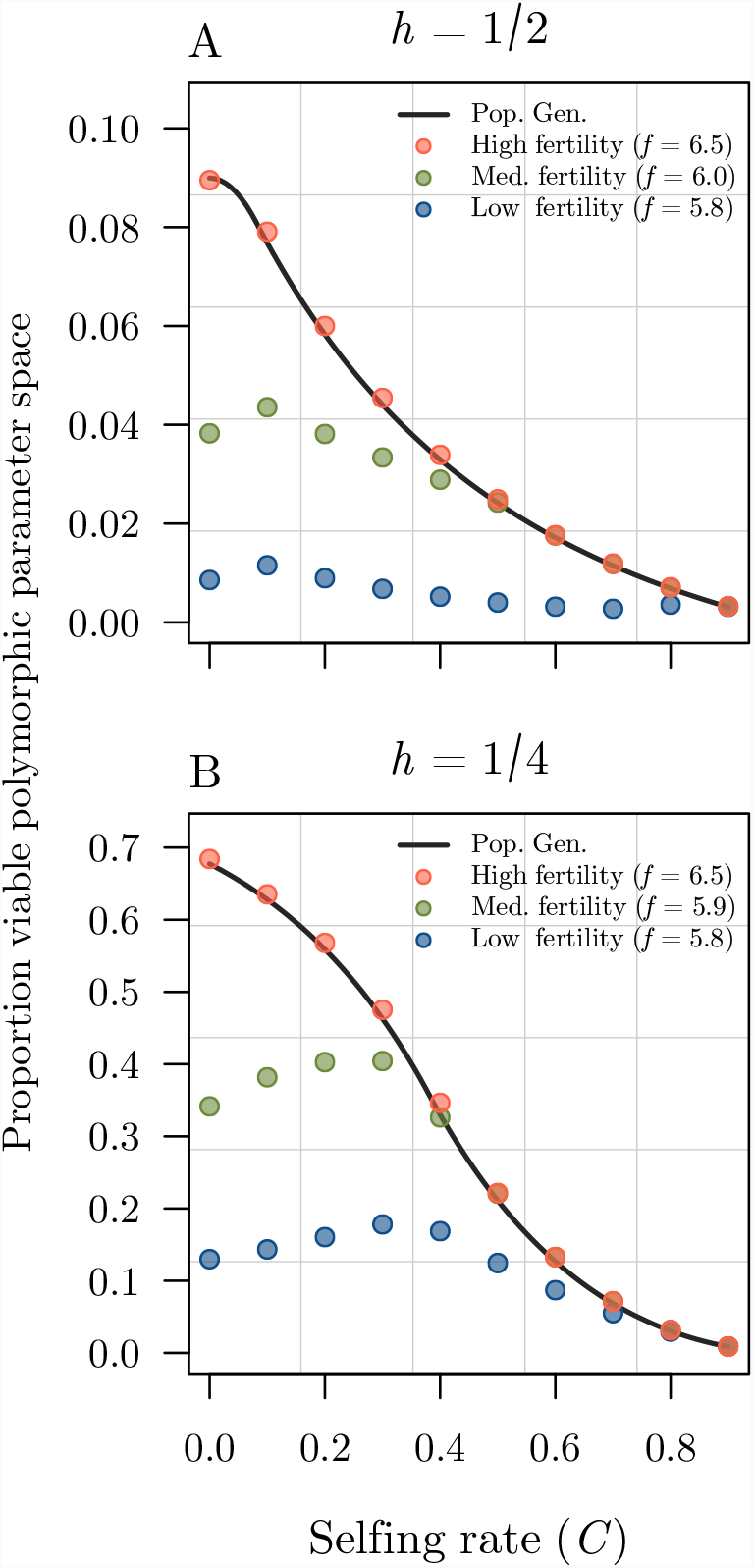
Proportion of demographically viable parameter space (out of total *s*_*f*_ × *s*_*m*_ space with max(*s*) = 0.15) in the absence of inbreeding depression (i.e., assuming *δ* = *δ*_*i*_ = 0, where *i* ∈ {*j, a, γ*}), plotted as a function of the population selfing rate. Results are shown for three fertility values corresponding to low, medium, and high fertility (blue, green, and red points respectively) under additive (*h* = 1/2; panel A), and partially recessive (*h* = 1/4; panel B) SA fitness effects. Each point was calculated by numerical integration of the corresponding SA invasion conditions and extinction threshold predicted by the mendelian matrix model (see Analyses section), while solid lines were produced by numerically integrating the analytic expressions for the single-locus invasion conditions from the population genetic models of Jordan and Connallon (2014) and Olito (2017) (solid black lines).

Invasion conditions for SA alleles in the evolutionary demographic model closely match the predictions from population genetic models (Jordan and Connallon, 2014; Kidwell et al., 1977; Olito, 2017). In particular, the demographic model recovers the classic “funnel-shaped” region of polymorphic *s*_*f*_ × *s*_*m*_ parameter space. The effects of the population selfing rate (*C*) and dominance (*h*) on SA polymorphism are also similar: self-fertilization (*i*) reduces the parameter space in which male-benefit alleles can invade, thereby increasing opportunity for spread of female-benefit alleles (e.g., contrast figure 1A with figure 1C); and (*ii*) dominance reversals (where deleterious SA fitness effects are partially recessive in each sex; *h* = 1/4) are much more permissive of SA polymorphism (e.g., contrast figures 1A-C with 1D-F) (Jordan and Connallon, 2014; Olito, 2017).

However, a key prediction from the evolutionary demographic model is that large fractions of SA polymorphic parameter space can be demographically inviable (fig. 1). The location of the extinction threshold, where the population intrinsic growth rate *λ*, = 1, is primarily determined by the fertility parameter (*f*) but is also influenced by the population selfing rate (*C*) and dominance of the SA alleles (*h*).

In populations with high fertility (larger *f*), the proportion of demographically viable polymorphic parameter space converges on the predictions for total SA polymorphic space in population genetic models (fig. 2). In obligately outcrossing populations (including dioecious/gonochoristic populations; where *C* = 0), lower fertility can result in a significant reduction of demographically viable polymorphic parameter space. The effect is weaker in populations with intermediate selfing rates (compare *C* = 0 vs. *C* > 0) because self-fertilization generates a greater proportion of offspring that are homozygous for the female-beneficial allele relative to heterozygotes. In other words, self-fertilization reduces the opportunity for selection favouring the female-deleterious allele, thereby reducing the equilibrium load on female fecundity and allowing partially selfing populations to remain viable under selection inten-sities that would cause extinction in an outcrossing population. The combination of protection from reduced female fertility and reduced total polymorphic parameter space caused by selfing results in populations with intermediate selfing rates having the greatest proportion of demographically viable parameter space at medium and low fertilities (fig. 2, med. and low. fertility values).

### Demographic effects of inbreeding depression

Unlike previous population genetic models, which assume constant population sizes (Jordan and Connallon, 2014; Olito, 2017), mortality caused by inbreeding depression can strongly influence population persistence in our evolutionary demographic model. Populations with high fertility rates can maintain positive population growth rates despite this additional mortality. This causes a greater proportion of SA polymorphic parameter space to be demographically viable, with the demographic model predictions converging on those from population genetic models in high-fertility populations (fig. 3). For populations with lower fertility rates, the proportion of demographically viable polymorphic parameter space matches the population genetic model predictions under complete outcrossing; however, as the selfing rate increases, demographic viability eventually crashes when the population can no longer sustain the concomitant increase in mortality due to inbreeding depression (fig. 3).

**Figure 3:**
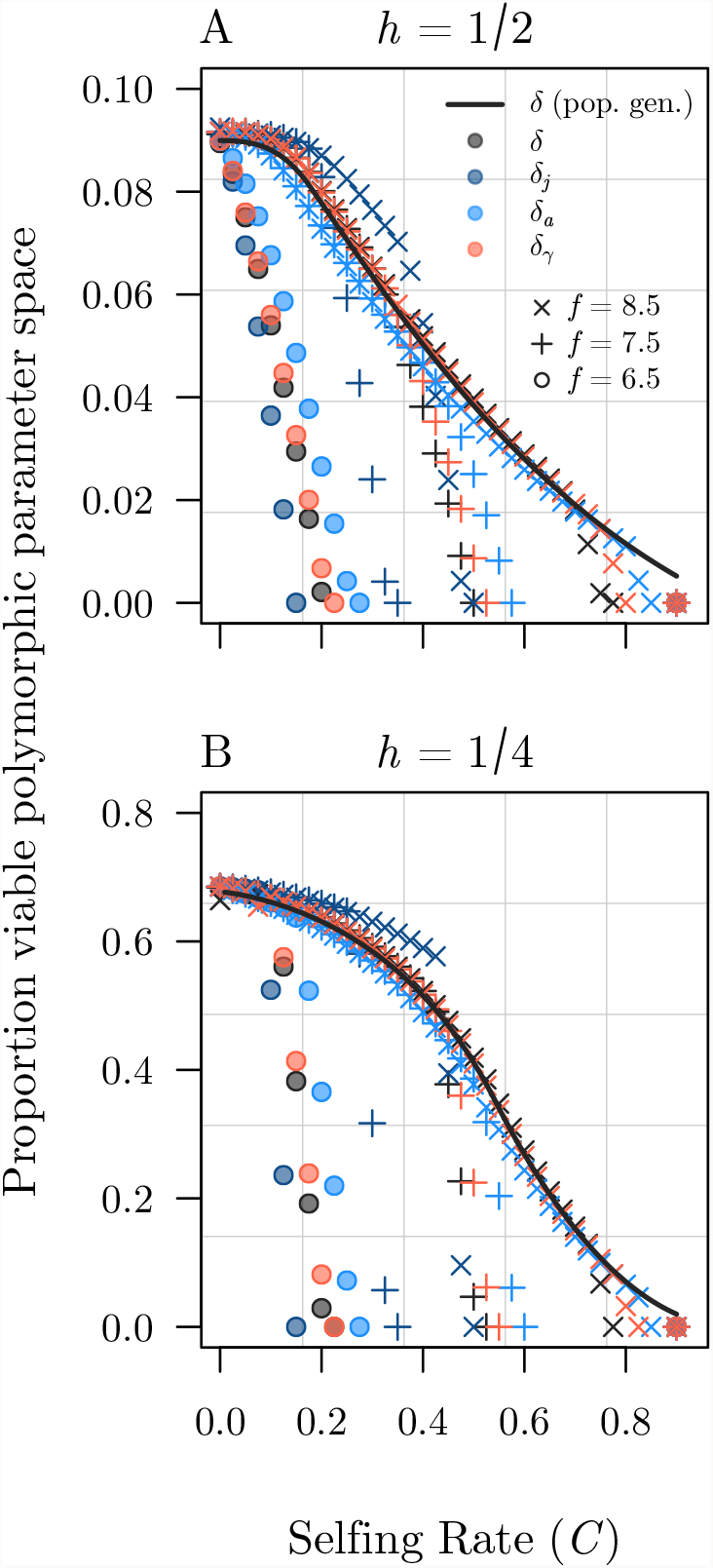
Effects of early- and late-acting inbreeding depression on the proportion of demographically viable parameter space (out of total *s*_*f*_ × *s*_*m*_ space with max(*s*) = 0.15), plotted as a function of the population selfing rate. In all plots, the strength of inbreeding depression decreases as the selfing rate goes up following a simple model of purging recessive deleterious mutations (see Analyses). Only single inbreeding depression terms (*δ* and *δ*_*i*_, where *i* ∈ {*j, a, γ*}, indicated in the legend) are allowed to vary at one time (all others are set to 0). Results are shown for three fertility values (*f* = {6.5, 7.5, 8.5}) under additive (*h* = 1/2; panel A) and partially recessive (*h* = 1/4; panel B) SA fitness effects. Each point was calculated by numerically integrating the corresponding SA invasion conditions and extinction threshold predicted by the model (see Analyses section), while solid lines were produced by numerically integrating the single-locus invasion conditions from the population genetic models of Jordan and Connallon (2014) and Olito (2017) (solid black lines).

Mortality from inbreeding depression quickly outweighs the beneficial effect of selfing on the equilbirium female load that was apparent in the previous section. In contrast to our earlier results, pre-dominantly outcrossing populations are predicted to have the highest proportion of demographically viable polymorphic space when inbreeding depression is taken into acount (compare fig. 2 with fig. 3). Regardless of the life-history stage at which inbreeding depression affects survival, populations with intermediate to high selfing rates are unlikely to harbour SA polymorphism unless they can afford the resulting loss of self-fertilized ovules/offspring. An interesting alternative interpetation of these results is that populations with intermediate to high selfing rates and population intrinsic growth rates near one are vulnerable to extinction if a sexually antagonistic allele invades the population.

The point in the life-cycle where inbreeding depression manifests can influence the threshold selfing rate at which demographically viable polymorphic parameter space crashes. Our results indicate that population viability was most sensitive to inbreeding depression affecting juvenile survival rates (*δ*_*j*_; fig. 3, dark blue points), while early-acting inbreeding depression (*δ*, ovule abortion shortly after fertilization) had a similar effect on population viability as late-acting inbreeding depression affecting adult survial (*δ*_*a*_) and juvenile-to-adult transition rates (*δ*_*γ*_). Inbreeding effects on juvenile survival had the strongest effect on population viability because on average individuals will spend multiple time steps in the juvenile stage before they mature. At each time, juveniles have a probability *γ* to mature and a probability *σ*_*j*_(1− *δ*_*j*_) to survive. Inbreeding depression at the juvenile stage therefore makes it harder to survive long enough to mature. Although early-acting inbreeding depression (*δ*) actually manifests earlier in the life-cycle than juvenile survival, it acts only once by influencing the total number of self-fertilized zygotes that become juveniles. Note, however, that the relative strength of inbreeding depression at different stages of the life history will depend on the interaction between the various inbreeding depression parameters, which we have precluded from our analyses.

### Case study: *M. guttatus*

As an illustrative example of how our model can be used to explore whether demographic rates observed in natural populations appear likely to support balanced SA polymorphisms, we parameterized the model using empirically estimated demographic rates and fitness data for natural populations of the hermaphroditic flowering plant *Mimulus guttatus* (Scrophulariaceae; now known as *Erythranthe guttata*). *M. guttatus* is an herbaceous, self-compatible wildflower native to western North America that exhibits remarkable among-population variation in numerous life-history and reproductive traits including selfing rates, inbreeding depression, floral morphology, and annual-to-perennial life-history (e.g., Ritland, 1990; Ritland and Ganders, 1987; Willis, 1993, 1999a,b; Wu et al., 2008). Moreover, detailed demographic studies have been conducted on multiple populations of *M. guttatus*, with demographic data available on the public demographic database COMPADRE (Plant Matrix Database, 2020). Below, we briefly outline how we parameterized our model using the available data; full details are provided in Appendix C.

#### Demographic and fitness data for M. guttatus

To parameterize our model we utilize extensive demographic data reported in a large-scale study of local adaptation using experimental populations of *M. guttatus* in Stanislaus National Forest (California, USA) in 2012 and 2013 (Peterson et al., 2016). We leverage their common-garden experimental design to focus on a comparison of the consequences of SA selection for two experimental populations with contrasting demographic rates. The first was a locally adapted ‘Eagle Meadows’ population (data from 2012), while the second was an experimental population composed of multiple non-locally adapted ‘low-elevation perennials’ (data from 2013). The vital rate estimates for the Eagle Meadows population are as follows: seed bank survival (*D* = 0.534), seed germination rate (*G* = 0.469), flower production (*F* = 0.64), ovules per flower (*O* = 614), seedling recruits proportional to clonal rosette recruits (*A* = 6.7 × 10 ^−4^), overwinter survival (*S* = 0.179), and rosette production (*R* = 8.71). The corresponding estimates for the low-elevation perrenials are: *G* = 0.652, *F* = 4.09, *O* = 494, *S* = 0, and *R* = 0 (see corrected Tables. 1 and S2 in Peterson et al. 2017). The same estimates for *D* and *A* were used for all populations. The resulting transition matrices for this population involved three life-history stages (*w* = 3; seed, seedling, and rosette), and individual elements of the transition matrix (**Ã**) were calculated as products of the above rates (see Matrix 1 in Peterson et al. 2016, also Eq(C1) in our Appendix C).

Estimates of selfing rates and inbreeding depression were not available for the same experimental populations, but are available for a variety of other western USA *M. guttatus* populations. Selfing rate estimates vary in magnitude from near complete outcrossing to predominant selfing (*C* ≈ 0 to 0.75; Ritland 1990; Ritland and Ganders 1987; Willis 1999b). Estimates of inbreeding depression at several of the life-history stages/fitness components that were included in the data of Peterson et al. (2016) are available for two intensively studied populations in the Cascade Mountains of Oregon (Iron Mountain and Cone Peak; Willis 1993, 1999a,b). Using the data provided in Willis (1993), we estimated the proportional decrease due to inbreeding depression in seed germination rate (*δ*_*G*_ = 0.085), flower number (*δ*_*F*_ = 0.2), and overwinter survival (*δ*_*S*_ = 0.38). The largest field-estimated selfing rate for the Iron Mountain population was *C* = 0.29 (Willis, 1993).

Using these combined demographic rates, selfing rates, and inbreeding depression estimates, we constructed a corresponding stage × genotype mendelian matrix model with a single SA locus affecting female and male fertility (as described above). With the empirically parameterized model, we are able to make predictions about the genetic and demographic outcome of SA selection a single locus in hypothetical populations with the same demographic rates as observed in Peterson et al. (2016), for a range of selfing and inbreeding depression rates observed in other natural populations. We stress, however, that these are illustrative rather than explicit predictions of the likelihood of SA polymorphism in any specific population, and they ignore measurement error for the estimated demographic rates.

Interestingly, a polymorphic chromosomal inversion (inv6) with apparently SA fitness effects has been identified in the Iron Mountain population of *M. guttatus* (Lee et al., 2016). inv6 segregates at moderate frequency (about 8%), and carriers suffer an approximately 30% loss in pollen viability, but also increased flower (and therefore ovule and pollen) production. The genetic basis of these effects remain unclear – though they likely polygenic – but the net result is a “supergene” with remarkably strong effects on both female and male fertility that segregates as a single diallelic locus. We estimated selection coefficients for the effect of inv6 on pollen production (i.e., taking into account the simultaneous effect on flower number) and ovule production from the data reported in Lee et al. (2016) under the relatively conservative assumptions that pollen/ovule production is proportional to flower number, and additive SA fitness effects (dominance coefficients could not be estimated from field data; see Appendix C for details of estimating selection coefficients). Under these assumptions, the average selection coefficient for inv6 on flower production across two years in the field was (*s*_*f*_, *s*_*m*_)= (0.3, 0.31). As a final proof-of-concept for our empirically parameterized model (e.g., Servedio et al., 2014), we asked whether, given the above biologically grounded fitness effect estimates, inv6 appears to fall in demographically viable SA polymorphic parameter space (see fig. 4), as might be expected given its observed frequency in the Iron Mountain population.

**Figure 4:**
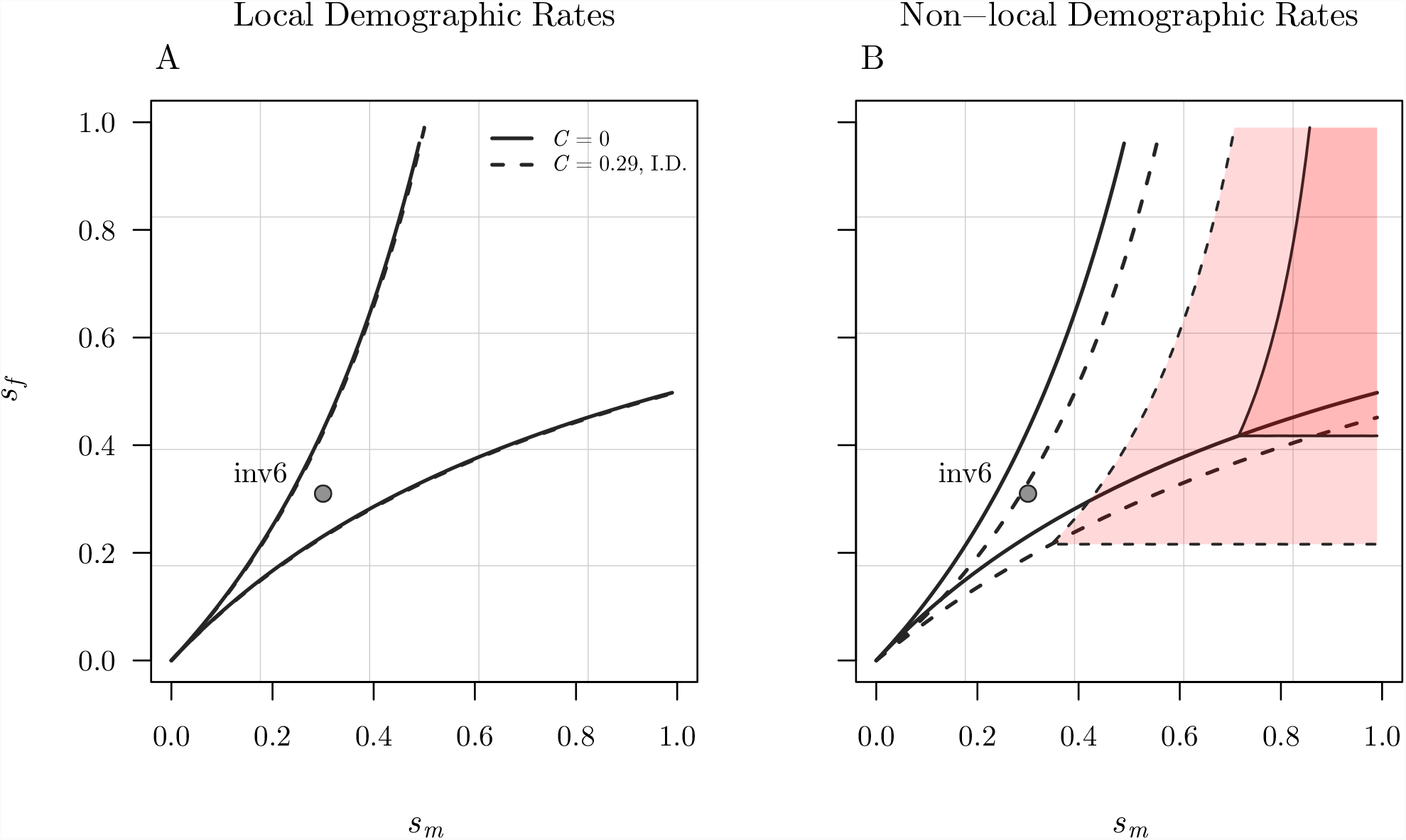
Illustration of model predictions using empirically estimated demographic rates for *M. guttatus*. Results are shown for two hypothetical populations using demographic rates for locally adapted (Eagle Meadows; panel A), and non-local (Low-Elevation Perennial; panel B) populations reported by Peterson et al. (2016). Invasion conditions and extinction thresholds were calculated using the conservative assumption of additive fitness effects in both sexes (*h*_*f*_ = *h*_*m*_ = 0.5) for two parameter conditions: obligate outcrossing (solid black lines, dark red shaded region) and partial selfing with inbreeding depression using the highest field-estimate of selfing, and inbreeding depression parameters (*δ*_*i*_) calculated for the Iron Mountain population of Willis (1993) (dashed lines, light red shaded region). The location of inv6 is also shown on both plots, using selection coefficients calculated from field estimates of male and female fitness components from Lee et al. (2016) under the relatively conservative assumption of additive fitness effects in both sexes, where (*s*_*f*_, *s*_*m*_) = (0.30, 0.31). Note that in panel A the invasion conditions for partial selfing with inbreeding depression (dotted lines) are nearly indistinguishable from those for obligate outcrossing (solid lines).

#### Polymorphism in M. guttatus

The Eagle Meadows and Low-Elevation Perennial populations of Peterson et al. (2016) had contrasting demographic rates that strongly affected the scope for demographically viable SA polymorphism. The Eagle Meadows population was locally adapted with a very high intrinsic growth rate (*λ* ≈ 1.7). This growth rate was sufficiently high that all *s* _*f*_ × *s*_*m*_ selection parameter space (where *s*_*f*_, *s*_*m*_ ∈ (0, 1]) remained demographically viable, regardless of the selfing rate and effects of inbreeding depression (*C* = 0 or 0.29; fig. 4A). In contrast, the Low-Elevation Perennial population had a much lower, but still positive, intrinsic growth rate (*λ* ≈1.08). Due to the slower growth rate, not all of the *s*_*f*_ × *s*_*m*_ selection parameter space was demographically viable: extinction thresholds appear under both complete outcrossing and partial selfing with inbreeding depression (fig. 4B). The different demographic rates from these two populations also resulted in slightly different invasion conditions when inbreeding depression was taken into account (compare dashed with solid lines in fig. 4A and B).

When using demographic rates for either the Eagle Meadows or Low-Elevation Perennial popula-tions, inv6 falls within demographically viable polymorphic parameter space (fig. 4). Whether under obligate outcrossing or partial selfing, inv6 always falls squarely in the middle of SA polymorphic space when using the locally adapted Eagle Meadows demographic rates. In contrast, when the selfing rate is at the higher end of empirical estimates for the Iron Mountain population in which inv6 has been documented (*C* = 0.29), inv6 falls nearer the upper boundary but still within SA polymorhic space when using the Low-Elevation Perennial demographic rates. This happens because under the empirical estimates of selfing and inbreeding depression, the polymorphic parameter space shifts downwards. Additionally, inv6 falls much closer to the extinction threshold under partial selfing, suggesting that even relatively small perturbations to demographic rates or selection coefficients could result in non-locally adapted populations being unable to support the demographic costs associated with segregating SA alleles with selection coefficients of similar magnitude to inv6.

## Discussion

Classic population genetics theory predicts that sexually antagonistic selection is unlikely to maintain genetic variation except under narrow conditions, with polymorphism requiring either finely balanced or unusually strong selection, or partially recessive fitness effects through each sex (Connallon and Clark, 2014; Kidwell et al., 1977; Pamilo, 1979; Prout, 2000). Extensions of the theory have identified numerous ways in which the conditions for polymorphism become more permissive in both dioecious and hermaphroditic organisms, including genetic linkage of SA loci, the evolution of sex-specific dominance, population subdivision, and life-cycle complexity (e.g., Connallon et al., 2019; Jordan and Charlesworth, 2012; Jordan and Connallon, 2014; Olito et al., 2018; Patten et al., 2010; Spencer and Priest, 2016). However, by ignoring the demographic consequences of SA genetic variation, these population genetic models have missed the possibility that SA polymorphisms may not be viable under realistic parameter conditions, and therefore unlikely to be observed in natural populations. By linking the individual-level fitness consequences of SA selection to population level consequences, our theoretical framework provides several key insights into the processes shaping SA genetic variation in natural populations.

The first and central finding of our study is that when intrinsic population growth rates approach one, the deleterious effects of segregating male-beneficial SA alleles on female fecundity can result in extinction over much of the parameter space where SA polymorphism is maintained. Since intrinsic growth rates far exceeding one suggest rapid exponential growth, they are generally rare (with the notable exception of recently introduced, invasive populations), suggesting that our model predictions may be highly relevant for SA polymorphism in many real-world populations. In addition, we find that much of the parameter space where a male-beneficial allele fixates is demographically inviable for populations whose intrinsic growth rates are close to one prior to invasion. This demographic consequence of masculinization is rarely considered (Hitchcock and Gardner, 2020). Moreover, these findings complement recent theoretical and empirical studies indicating that SA selection is likely to be both condition dependent, and stronger in locally-adapted populations near the center of a species’ range, where population growth rates are expected to be high (Berger et al., 2014; Connallon, 2015).

We also find that demographically viable parameter space is often biased towards alleles with stronger selection through the female than male sex function. That is, given natural variation in population growth rates, the most demographically viable (and therefore observable in natural populations) outcomes of SA selection are either fixation of a female-beneficial allele, or polymorphisms involving low-frequency female-deleterious alleles. This key prediction is supported by a series of experimental results in seed beetles (*Callosobruchus maculatus*) which show that male-beneficial SA genotypes are less likely to contribute positively to population growth rates, and are more susceptible to extinction under environmental stress or inbreeding (Berger et al., 2014, 2016; Grieshop et al., 2017). An interesting corrollary of our findings is that, since strong SA fitness effects often lead to extinction, the observable SA genetic variation in natural populations may often be under weak selection, and therefore strongly susceptible to genetic drift whether or not selection favours the fixation of one allele or balancing selection (Connallon and Clark, 2012).

In hermaphroditic populations, self-fertilization can alleviate the demographic costs of balanced SA polymorphisms under some conditions, however, the concommitant effects of inbreeding depression generally exacerbate them in populations with mixed-mating systems. This prediction is in stark contrast to previous population genetics models of SA selection in hermaphrodites, where the sole effect of inbreeding depression is to reduce the population effective selfing rate through the loss of selfed-zygotes, thereby expanding polymorphic parameter space (Jordan and Connallon, 2014; Olito, 2017). In our model, this reduction in effective selfing is accompanied by significant mortality due to inbreeding depression (whether early- or late-acting), which can quickly tip partially-selfing populations over the brink to extinction. Beyond the maintenance of SA polymorphisms, this finding underscores a simple but important point that is often overlooked in studies of the evolution of self-fertilization and selfing-syndromes, which tend to emphasize the coevolution of the deleterious mutation load and mating system (e.g., Charlesworth and Charlesworth, 1987; Goodwillie et al., 2005; Lande and Schemske, 1985): highly fertile populations can better afford the severe demographic costs of inbreeding. This suggests that traits related to female fecundity, such as ovule and flower production, may strongly influence the distribution of successful transitions to self-fertilization among hermaphroditic taxa, as well as a variety of ecological correlates of selfing and mixed-mating (Goodwillie et al., 2005; Grossenbacher et al., 2015; Igic and Kohn, 2006).

Despite the demographic pitfalls associated with SA alleles, our example using *M. guttatus* appears to show that demographic rates observed in some real populations are capable of sustaining large regions of viable SA polymorphic space. The example also appears to provide some empirical support for the conjecure that locally-adapted populations are more likely to harbor SA polymorphisms than marginal or non-locally adapted ones; inviable polymorphic parameter space only occured when using the demographic data for non-local high-elevation perennial populations. Although we cannot make concrete predictions for inv6 in the Iron Mountain population in which it was observed, it is interesting that the estimated SA fitness effects place this polymophic inversion squarely in demographically viable polymorphic parameter space predicted by our model. The available data do not allow for confident estimation of selection coefficients for inv6 (and even require making assumptions about dominance), yet our theoretical predictions are encouragingly consistent with the available data that inv6 is segregating at intermediate frequencies in the large and locally adapted Iron Mountain population.

Overall, our findings provide a more nuanced picture of the nature of SA genetic variation that we should expect to find in natural populations, where the fate of SA alleles and the populations harboring them is determined jointly by evolutionary and demographic processes.

### Extensions and future directions

By combining the tools of demography and population genetics, the framework we present here enables the exploration of interactions between life cycle complexity, mating system, and sexual antagonism. For the sake of simplicity, we made a variety of simplifying assumptions in our analyses, and a variety of extensions are possible. For example, we used a simple life cycle with just two stages for most of this paper, adults and juveniles (the *Mimulus* example has 3 stages). However, it is possible to include additional age classes, allowing us to explore whether the scope for SA selection to maintain polymorphisms is affected by whether a species exhibits positive or negative senescence (Jones et al., 2014). Put another way, does the shape of the survival curve affect the demographic and population genetic consequences of SA selection, and how does this interact with the age at which a sexually antagonistic allele is expressed? What if the allele not only affects male and female fertility but also affects survival of individuals? Additionally, how would the scope for SA polymorphism be affected by density-dependence acting on different life history stages?

Like most demographic matrix models, ours also assumes female demographic dominance, where population growth rates are determined entirely by female fecundity (Caswell, 2001; Iannelli et al., 2005; Pollard, 1975). Yet, many hermaphroditic populations experience limitation of reproductive success due to both quantity and quality of male gametes (e.g., Aizen and Harder, 2007; Harder et al., 2016; Yund, 2000). Explicitly modeling male gamete production is a natural extension to our modeling framework, and would enable us to analyze how tension between the demographic consequences of SA fitness variation through both female and male function alters or abolishes asymmetries in the extinction thresholds and polymorphic parameter space (e.g., Tazzyman and Abbott, 2015). Interestingly, different forms of self-fertilization can aid in reproductive assurance, and should therefore have demographic consequences – for example, under pollen limitation and delayed selfing (where only ovules that fail to receive outcross pollen are selfed), the selfing rate will be a function of genotype frequencies because these will directly influence pollen production (Harder and Barrett, 2006).

Another major simplifying assumption in our model was that how individuals themselves were produced (i.e., by selfing or outcrossing) affects the level of inbreeding depression they suffer. In reality, the history of consecutive generations of inbreeding in each individual’s lineage will influence the severity of inbreeding depression they experience, particularly when inbreeding depression is caused primarily by recessive deleterious mutations. It would be an interesting and feasible extension of our modeling framework to expand the individual state space to include selfing cohorts (i.e., first generation selfing, second generation selfing, etc.), as in the models of Kelly (1999, 2007), thereby enabling a more biologically plausible approach to modelling self-fertilization.

Grieshop et al. (2017) found that genotypes with sexually antagonstic alleles that are male beneficial experience higher levels of inbreeding depression than genotypes with female beneficial SA alleles. This effect can easily be included in our model by making the inbreeding depression parameters, *δ* and *δ*_*i*_, a function of the genotype of the individual. Such an effect would further reduce the demographically viable polymorphic parameter space, and increase the bias in viable SA polymorphisms towards alleles with weaker selection in females.

## Conclusion

Despite a surge of interest in eco-evolutionary dynamics, the demographic consequences of intralocus sexual antagonism have rarely been modeled (but see Harts et al. 2014; Kokko and Brooks 2003; Matthews et al. 2019). In contrast, models of the population dynamical consequences of interlocus sexual conflict are more common (e.g., Martínez-Ruiz and Knell, 2017; Tanaka, 1996). We found that including basic demography can have a significant impact on traditional population genetic results, as has been suggested previously both by theoretical and empirical work (Berger et al., 2016; Grieshop et al., 2017; Kokko and Brooks, 2003). Although the potential negative consequences of sexual conflict for population viability have been known for some time, this aspect of sexual antagonism has rarely been considered in models investigating the scope for intralocus sexual antagonism to maintain genetic variation.

Demographic models connect individual level traits to population level consequences. As a consequence of their focus on individuals, demographic models are ideal for linking to experimental or field data, as demonstrated with our *Mimulus* case study. Doing so allows the field of population genetics to move from fitness as an abstract scalar metric towards the fitness of an entire life cycle as calculated from observed rates of age- or stage-specific survival and fecundity rates.

## Appendix A

### Population projection matrix

The complete population projection matrix **Ã** population vectors: consists of 3 × 3 blocks, which act on the genotype specific population vectors:

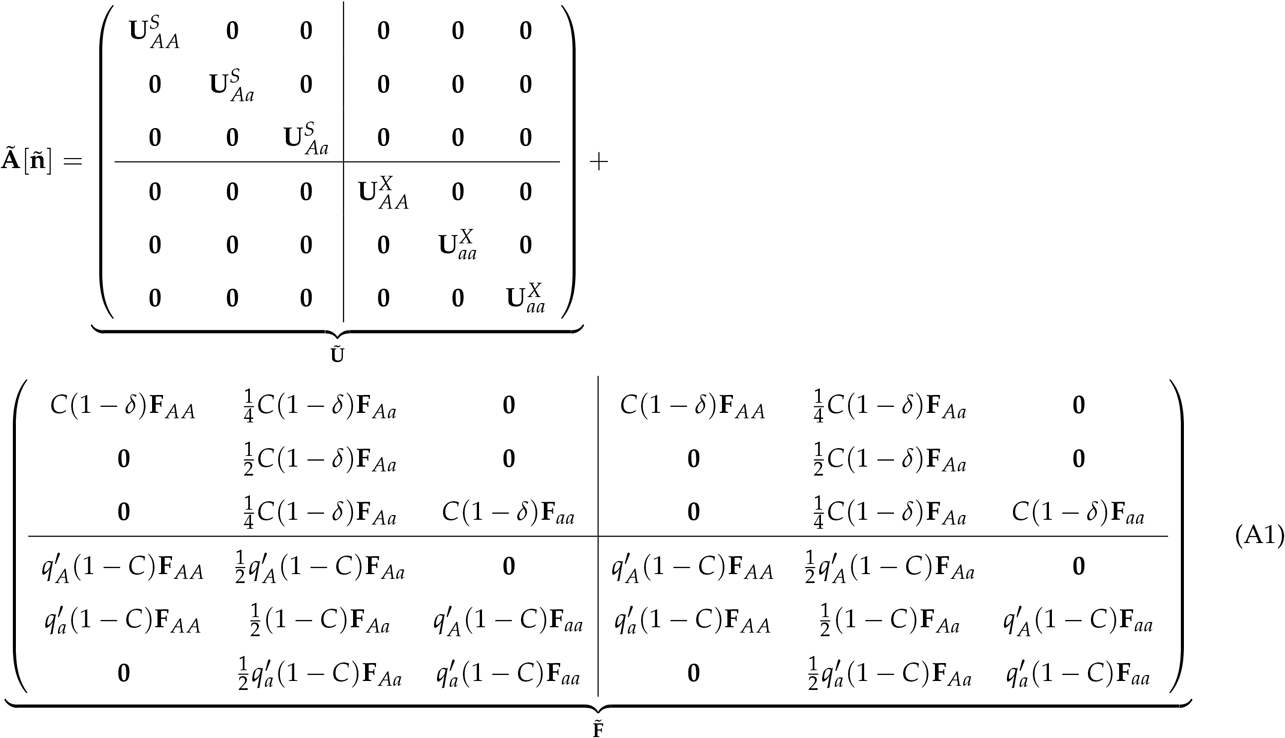

with symbols as defined in the main text. The survival matrices appear on the diagonal because individuals do not change their genotype once they are born. The fertility matrix incorporates the process of Mendelian inheritance and is an extension of the fertility matrix derived by de Vries and Caswell (2019a).

The first block column of **Ã** describes the production of offspring by an *AA* female with stage-specific fertility rates **F**_*AA*_ by both selfing and outcrossing. The probability of picking an *A* allele out of the pool of available male gametes. When reproducing by selfing, this is entirely determined by the probability of sampling an *A* allele after Mendelian segregation. For outcross reproduction the probability, and hence the probability of this *AA* female producing an *AA* offspring, is 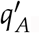, as derived above. Conversely, the probability of picking an *a* allele and producing an *Aa* offspring is zero for an *AA* female when selfing, but 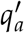 for outcrossing. Similarly, the middle column of block matrices are offspring produced by *Aa* females, which can produce offspring of all 3 genotypes.

A full derivation of the model, including all component matrices, is provided in the Online Supplementary Material.

## Appendix B

### Eigenvalues for invasion analysis

The invasion analysis is made easier if we first reorder the population vector by genotype, then by how individuals were produced (selfing vs. outcrossing), and finally by stage. Ordering by genotype first facilitates the invasion analysis in two ways. First, the resulting Jacobian matrix is upper block triangular, and the eigenvalues of **M** are therefore the eigenvalues of the diagonal blocks. Second, the blocks along the diagonal correspond to perturbations in each of the the three genotype ‘directions’ at the boundary equilibria. As outlined in the Online Supplementary Material, the stability of both boundary equilibria 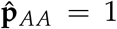 and 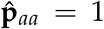 are determined by the central block of the Jacobian, **M**_22_, which corresponds to perturbations in the direction of heterozygote *Aa* genotypes. A boundary equilibrium is unstable to invasion by the rare allele if the largest absolute value of the eigenvalues of the Jacobian matrix, the leading eigenvalue, evaluated at the equilibrium is greater than 1. The resulting conditions for a protected polymorphism require that

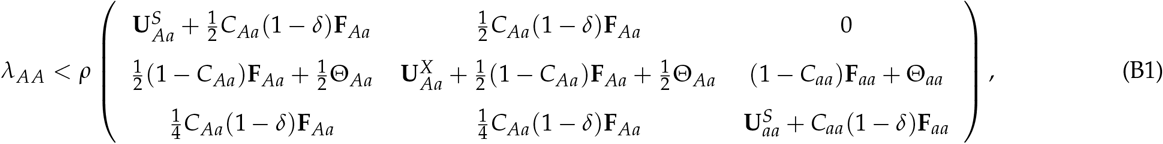

and

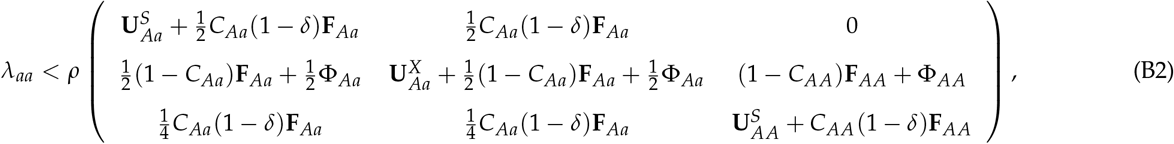

is satisfied, where where *ρ*(·) represents the spectral radius, and 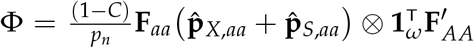. *ρ*(**M**_22_) gives the ergodic growth rate of perturbations of the boundary equilibrium according to the linear approximation, and *λ*_*i*_ is the intrinsic growth rate for the homozygous genotype being invaded at each boundary (*i* ∈ (*AA, aa*)).

In our numerical simulations, equations Eq(B1) and Eq(B2) were evaluated to determine the invasion thresholds for the relvant parameter conditions described in each figure.

Under obligate outcrossing (when *C* = 0), the leading eigenvalue evaluated at the 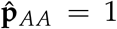 and 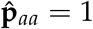 boundaries is equal to

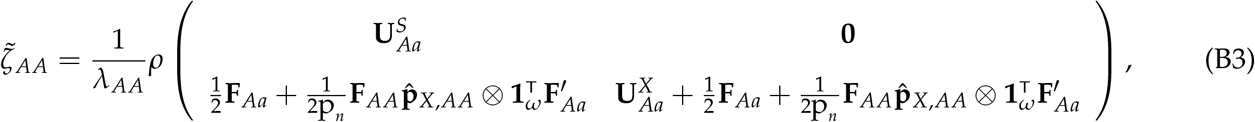

and

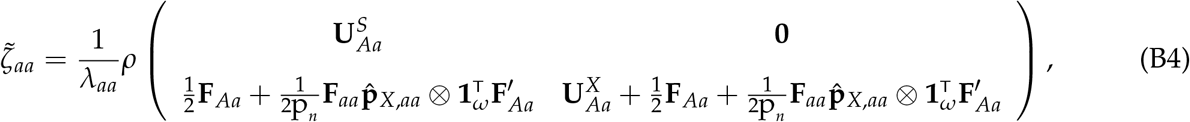

which reduce to

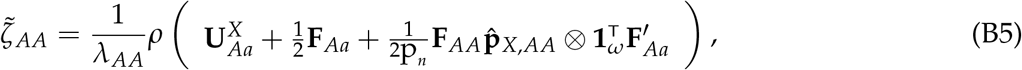

and

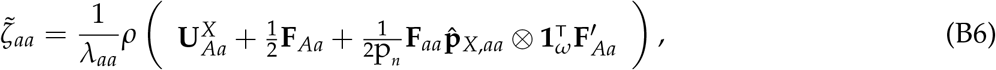

when 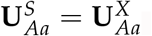. These eigenvalues are identical to those derived by de Vries and Caswell (2019a), confirming that our model predictions reduce to the two-sex model under obligate outcrossing, as expected.

## Appendix C

### Demographic and empirical data for *M. guttatus* case study

#### Demographic data

We used demographic data for multiple experimental populations of *M. guttatus* from a large-scale common garden experiment conducted in Stanislaus National Forest (California, USA) in 2012 and 2013 (Peterson et al., 2016). Specifically, we used data from two experimental populations with contrasting demographic rates: the locally adapted ‘Eagle Meadows’ population (data from 2012), and a non-local ‘low-elevation perennials’ population (data from 2013). The vital rates used in theirs, and our, calculations are summarized below in Table C1 (see corrected Tables. 1 and S2 in Peterson et al. (2016, 2017)). Note that the same estimates for seed bank survival (*D*) and Seedling recruitment (*A*) were used for all populations in the study.

The resulting transition matrices for these populations involve three life-history stages (*w* = 3; seed, seedling, and rosette; see Matrix 1 Peterson et al. 2016), and can represented as the sum of survival and fertilitiy matrices as follows:

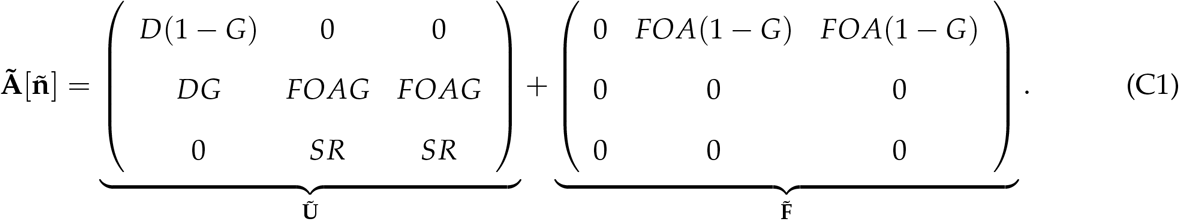

Overall transition matrix data for the same populations is also available on the public demographic database COMPADRE (Plant Matrix Database, 2020), but are not decomposed into the product of terms described in Eq(C1) above.

As described in the main text, we incorporated empirical estimates of the population selfing rate and inbreeding depression for two intensively studied populations in the Cascade Mountains of Oregon into our parameterized model (Iron Mountain and Cone Peak; Willis 1993, 1999a,b). Using the data provided in Table. 2 of Willis (1993), we calculated the proportional decrease due to inbreeding depression in three fitness components that were also included in vital rate estimates of Peterson et al. (2016): seed germination rate (*G*), flower number (*F*), and overwinter survival (*S*). In each case, we calculated the inbreeding depression terms as *δ*_*i*_ = 1 −*w*_inbred_/*w*_outcross_. To incorporate these terms into our evolutionary demographic model, we multiplied each of the three terms *G, F*, and *S* in Eq(C1) by a corresponding proportional inbreeding depression term (1 − *δ*_*i*_), where *i* ∈ {*G, F, S*}. The largest field-estimated selfing rate for this same Iron Mountain population was *C* = 0.29 (Willis, 1993), which we use in our calculations for fig. 4.

**Table C1:** Vital rates estimated for *M. gutattus* experimental populations by Peterson et al. (2016)

| Vital rate estimated | Eagle Meadows (2012) | Low-Elevation Perennials (2013) |
| --- | --- | --- |
| Seed bank survival (D) | 0.534 | 0.534 |
| Seed germination rate (G) | 0.469 | 0.652 |
| Flower production (F) | 0.64 | 4.09 |
| Ovules per flower (O) | 614 | 494 |
| Seedling recruits (A) | $6.7 \times 10^{-4}$ | $6.7 \times 10^{-4}$ |
| Overwinter survival (S) | 0.179 | 0 |
| Rosette production (R) | 8.71 | 0 |
| Population intrinsic growth rate ($\lambda$) | 1.71 | 1.08 |

#### Selection on inv6

Lee et al. (2016) describe a polymorphic chromosomal inversion segregating in the Iron Mountain population of *M. guttatus*. Importantly, the study reports fitness effects of inv6 on both male and female fitness components. inv6 appears to cause a significant decrease in pollen viability, while simultaneously causing an increase in flower number (and therefore ovule and pollen production) which varies among years. Unfortunately, it was not possible to estimate the dominance of the fitness effects of inv6 in the field because individuals homozygous for the inversion were quite rare (although they were viable in greenhouse conditions). We therefore made the relatively conservative assumption that the fitness effects of inv6 on pollen viability and flower number were both additive. Under this assumption, it is possible to estimate selection coefficients for proxies of female and male fertilities.

We used flower number as a proxy for ovule production and hence female fertility. To calculate selection on flower number, we first calculated the geometric mean flower number in the field for wild-type and inv6 heterozygotes across the two years for which data was available (2012 and 2013), extrapolated the expected flower number for inv6 homozygotes (table C2), and calculated the relative fertility of each genotype given that inv6 homozygotes have the highest flower number (*w*_*i*_, where *i* ∈ {*inv*6/*inv*6, *inv*6/*w*+, *w* + /*w*+}). Using the relative fertility expressions outlined in Table C4, we then solved for *s*_*f*_.

**Table C2:** Selection coefficient estimates for inv6.

| Geom. Mean (2012 & 2013) | Genotype |  |  |
| --- | --- | --- | --- |
|  | *inv6/inv6* | *inv6/w+* | *w+ /w+* |
| Flower #: | 4.63 | 3.91 | 3.19 |
| Flower # × Pollen viab.: | 1.89 | 2.29 | 2.70 |
| Rel. Flower #: | 1 | 0.85 | 0.69 |
| Rel. Flower # × Pollen viab.: | 0.70 | 0.85 | 1 |

Table C3: Note that Values for *inv*6/*inv*6 homozygotes are extrapolated based on the assumption of additive effects of the inversion on flower and pollen production.

We used pollen production as a proxy for male fertility. We calculated average pollen production as the product of the geometric mean flower number and field estimated pollen viability for wild-type and inv6 heterozygotes, and extrapolated the expected pollen production for inv6 homozygotes (table C2). As before, we calculated relative male fertility 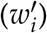 for each genotype given that wild-type homozygotes had the highest pollen viability, and used the expressions for 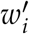 from Table C4 to solve for *s*_*m*_.

**Table C4:** Relative fertilities for inv6 genotypes (*w*_*i*_)

|  | Genotype |  |  |
| --- | --- | --- | --- |
|  | *inv6/inv6* | *inv6/w+* | *w+/w+* |
| Female function ($w_i$): | 1 | $1 - s_f/2$ | $1 - s_f$ |
| Male function ($w'_i$): | $1 - s_m$ | $1 - s_m/2$ | 1 |

Lee et al. (2016) also report greenhouse pollen viabilty data for several mapping populations of *M. guttatus*, which includes data for inv6 homozygotes, from which relative fertilities can be estimated. We chose to use the field estimated pollen viability and flower number data, despite the lack of inv6 homozygotes, because it probably gives a more accurate reflection of the fitness effects of inv6 in the field. Moreover, both approaches require making assumptions regarding the dominance effects of inv6. Given the obvious pitfalls of estimating selection on inv6 from the available data, it goes without saying that the location of inv6 in demographically viable polymorphic parameter space is a highly speculative, but nevertheless interesting, proof of concept for our model predictions.

## 1 Tables

Tables moved up to Model section in the main text.

## 2 Figure legends

Figure legends provided beneath each figure in the main text.

## Notes

https://anonymous.4open.science/r/SA-Hermaphrodites-wDemography-BF66

### Competing Interest Statement

The authors have declared no competing interest.

